# Aβ-Overlapping Ectodomain Binding of the Clinical-Stage TREM2 Agonist VG-3927

**DOI:** 10.64898/2026.03.02.709194

**Authors:** Sungwoo Cho, Moustafa T. Gabr

**Affiliations:** Department of Radiology, Molecular Imaging Innovations Institute (MI3), Weill Cornell Medicine, New York, NY 10065, USA

**Keywords:** TREM2, VG-3927, amyloid-β, Alzheimer’s disease, microscale thermophoresis

## Abstract

Triggering receptor expressed on myeloid cells 2 (TREM2) is a microglial immune receptor genetically and functionally linked to Alzheimer’s disease (AD). VG-3927, the first clinical-stage small-molecule TREM2 agonist, has been proposed to function as a transmembrane molecular glue and positive allosteric modulator (PAM). Whether it directly engages the extracellular ligand-recognition surface of TREM2 remains unknown. Here, we used a deep learning-based blind docking algorithm to map potential VG-3927 binding sites across TREM2 and identified a binding site within the ectodomain hydrophobic groove, a ligand-recognition surface previously implicated in Aβ and apoE binding. Microscale thermophoresis (MST) confirmed direct interaction of VG-3927 with TREM2 under optimized PEG-400 buffer conditions and independently demonstrated binding of Aβ^1-42^ to the receptor. Co-incubation with Aβ reduced the VG-3927 thermophoretic response, consistent with interference at an overlapping ectodomain binding surface. Consistently, Aβ induced a rightward shift in the VG-3927 dose–response curve in a Jurkat TREM2–DAP12 NFAT reporter assay and attenuated VG-3927-induced phospho-SYK signaling. Together, these findings support the presence of a previously unrecognized ectodomain interaction mode for VG-3927 and suggest that amyloid-associated ligand occupancy may modulate TREM2 agonist activity within the AD microenvironment.

## 1. Introduction

Triggering receptor expressed on myeloid cells 2 (TREM2) is a type I transmembrane glycoprotein predominantly expressed on microglia in the central nervous system (CNS).^1, 2^ TREM2 signals through its obligate adaptor protein DNAX-activating protein of 12 kDa (DAP12), mediating critical microglial functions including phagocytosis, lipid metabolism, inflammatory regulation, and survival.^3-53–5^ Genome-wide association studies identified loss-of-function variants in TREM2, most notably R47H, as among the strongest genetic risk factors for late-onset Alzheimer’s disease (AD), conferring a risk comparable to a single APOE ε4 allele.^6, 7^

The TREM2 ectodomain adopts an immunoglobulin (Ig)-like V-type fold and engages diverse ligands including phospholipids, apolipoproteins (apoE, CLU/apoJ), TDP-43, and amyloid-β (Aβ) oligomers.^8-13^ Structural studies by Kober et al. and Sudom et al. identified two principal ligand-binding surfaces on the TREM2 ectodomain: a large hydrophobic site located at the distal end of the molecule (CDR1, CDR2, CDR3 loops), and an electropositive basic site on the midsection.^14, 15^ Recent biophysical mapping by Greven et al. demonstrated that TREM2 utilizes this hydrophobic site as a shared surface to engage multiple AD-relevant ligands including apoE, TDP-43, and oAβ42.^16^ Critically, co-crystallization of TREM2 with an Aβ(1– 8) peptide confirmed that Aβ binds directly to the hydrophobic groove near CDR1.^17^

Aβ binding to TREM2 promotes microglial clustering around amyloid plaques and facilitates phagocytic clearance, a protective mechanism impaired by AD-associated TREM2 variants.^11, 18, 19^ The soluble ectodomain of TREM2 (sTREM2) has also been shown to directly bind Aβ fibrils via a partially overlapping surface, inhibiting secondary nucleation and enhancing cellular uptake.^20^ These findings underscore the importance of the TREM2 ectodomain surfaces in amyloid biology.

VG-3927 is the first small-molecule TREM2 agonist to enter clinical development. Preclinical characterization demonstrated that VG-3927 promotes anti-inflammatory microglial activation, suppresses neurodegeneration biomarkers in human iPSC-derived tri-cultures, and reduces AD-associated neuropathology in humanized TREM2 amyloidosis mice (hTREM2–5xFAD).^21, 22^ Mutagenesis mapping studies indicated that VG-3927 activity is on-target, functioning as a molecular glue and positive allosteric modulator (PAM) that potentiates TREM2 signaling via coordinated receptor clustering.^21^ However, no prior study has experimentally demonstrated ectodomain engagement by a small-molecule TREM2 agonist.

Importantly, our group recently reported the discovery of C1, a structural analog of VG-3927 with improved pharmacokinetic properties and proposed a potential ectodomain binding mode for this chemotype based on docking analyses.^23^ This observation prompted us to investigate whether VG-3927 itself might also engage the TREM2 ectodomain, and if so, whether this interaction overlaps the known Aβ-binding surface.

In this study, we employed DiffDock-L to perform an unbiased search for VG-3927 binding sites across the entire TREM2 structure. We identified a binding site within the TREM2 ectodomain hydrophobic groove and tested its functional relevance using orthogonal biophysical and cell-based assays, including MST, a Jurkat TREM2–DAP12 NFAT reporter assay, and phospho-SYK western blotting. Our findings support the presence of an ectodomain interaction mode for VG-3927 and indicate that Aβ can interfere with this interaction, thereby attenuating downstream TREM2 signaling.

## 2. Results

### 2.1 DiffDock-L Predicts a VG-3927 Binding Site on the TREM2 Ectodomain

To comprehensively map VG-3927 binding sites on TREM2, we performed blind docking simulations using DiffDock-L, which employs a diffusion-based generative model to predict ligand binding poses without prior knowledge of the binding pocket.^24^ Analysis of the top-ranked poses identified a binding site within the hydrophobic groove of the TREM2 ectodomain **(Figure 1A)**. Although the developer of VG-3927 (Vigil Neuroscience) has proposed that VG-3927 acts as a molecular glue promoting TREM2 clustering, the precise atomic-level binding site has not been publicly disclosed. Our DiffDock-L simulations predict that, in addition to the proposed transmembrane molecular glue mechanism, VG-3927 also engages a pocket formed by residues in the CDR1, CDR2, and CDR3 loops—the well-characterized hydrophobic pharmacophore previously identified as the Aβ and apoE binding region.

**Figure 1.**
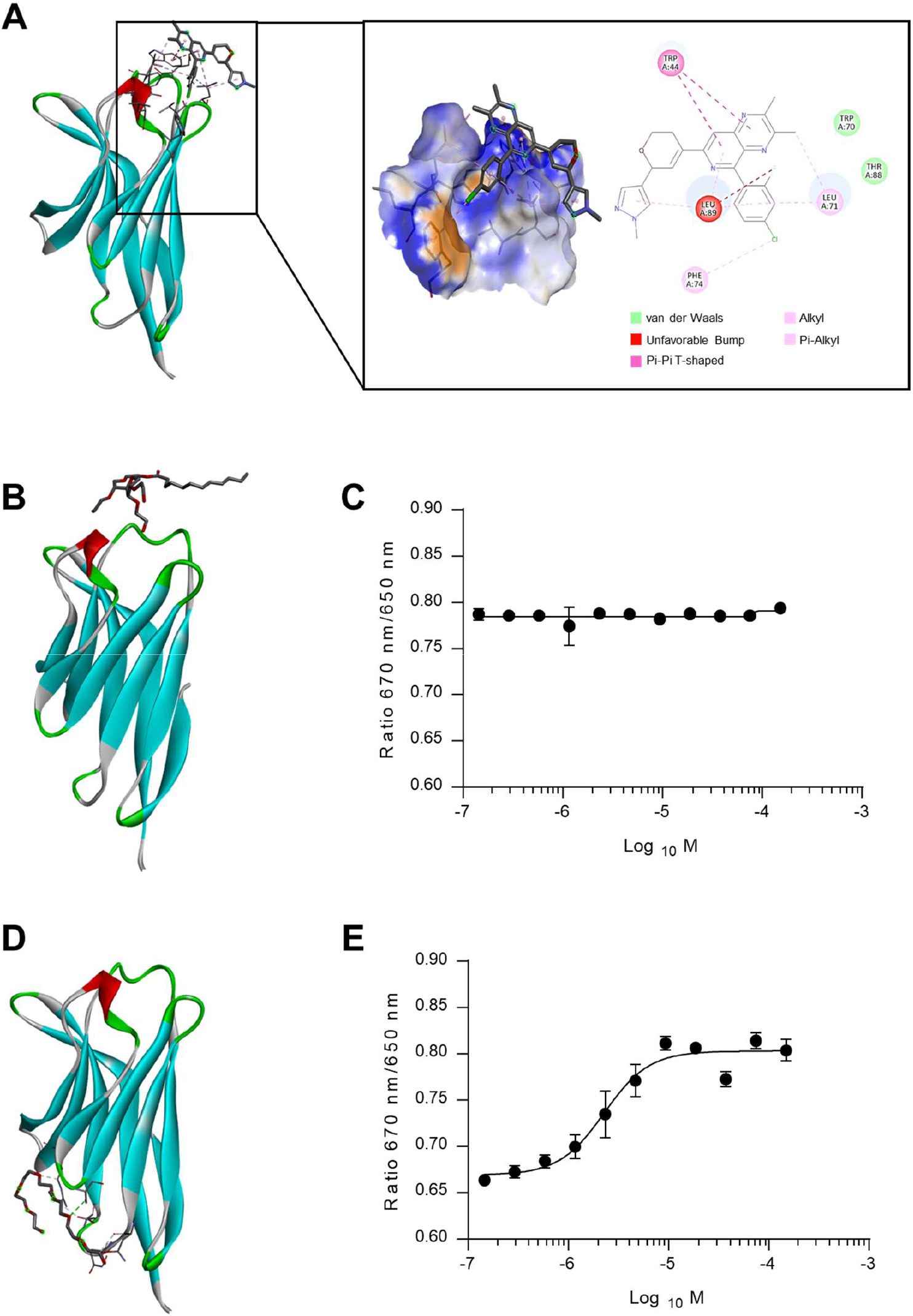
DiffDock-L identifies a novel VG-3927 binding site on the TREM2 ectodomain, and MST buffer optimization reveals detergent competition at the hydrophobic groove. (A) DiffDock-L blind docking results. Left: cartoon representation with VG-3927 (stick). Center: electrostatic surface representation. Right: 2D interaction diagram showing π–π T-shaped stacking (Trp44), van der Waals contacts (Trp70, Thr88), alkyl/π-alkyl interactions (Leu71, Phe74, Leu89), and an unfavorable steric contact (fluorine–Leu89). (B) Tween-20 occupies the TREM2 hydrophobic pocket. (C) MST in Tween-20 buffer: no VG-3927 binding detected. (D) PEG-400 does not occupy the hydrophobic pocket. (E) MST in PEG-400 buffer: VG-3927–TREM2 binding confirmed. Data are mean ± SEM, n = 3.

Detailed interaction analysis revealed that VG-3927 forms extensive hydrophobic contacts within this pocket **(Figure 1A, right panel)**. Specifically, the bipyridine core of VG-3927 engages in π–π T-shaped stacking with Trp44, while the 2-fluoro-4-chlorophenyl moiety forms alkyl and π-alkyl interactions with Leu71, Phe74, and Leu89. Additional van der Waals contacts are established with Trp70 and Thr88. Notably, an unfavorable steric bump was detected between the fluorine atom of VG-3927 and Leu89, suggesting that this position may be amenable to medicinal chemistry optimization.

### 2.2 Optimization of MST Conditions Reveals Detergent Interference at the Hydrophobic Site

To experimentally validate VG-3927 binding at the ectodomain, we employed microscale thermophoresis (MST). Initial experiments using standard buffer containing 0.05% Tween-20 failed to detect VG-3927–TREM2 interaction **(Figure 1B**,**C)**. We hypothesized that Tween-20, a non-ionic surfactant with significant hydrophobic character, might compete with VG-3927 for binding at the ectodomain hydrophobic pocket. Indeed, substitution with 0.05% PEG-400, a hydrophilic polyethylene glycol that does not occupy hydrophobic cavities, enabled robust detection of VG-3927–TREM2 binding **(Figure 1D**,**E)**. This detergent sensitivity provides indirect evidence that VG-3927 occupies the hydrophobic groove, consistent with the docking predictions and with the known hydrophobic character of this binding surface.

### 2.3 Aβ Interferes with VG-3927 Binding to TREM2

Having established that the proposed VG-3927 ectodomain binding site overlaps with the known Aβ interaction surface, we next examined whether Aβ could interfere with VG-3927 binding to TREM2. Using the optimized PEG-400 MST conditions, we first confirmed direct binding of Aβ_1–42_ to TREM2, with a measured Kd of 1.85 ± 0.24 μM **(Figure 2A)**, consistent with prior reports. We then performed competition experiments in which VG-3927 binding was assessed in the presence of 10 μM Aβ. Co-incubation with Aβ significantly reduced the VG-3927 MST signal **(Figure 2B)**, indicating that Aβ interferes with VG-3927 binding to TREM2 and supporting the idea that the two ligands share an overlapping binding surface within the TREM2 ectodomain hydrophobic site.

**Figure 2.**
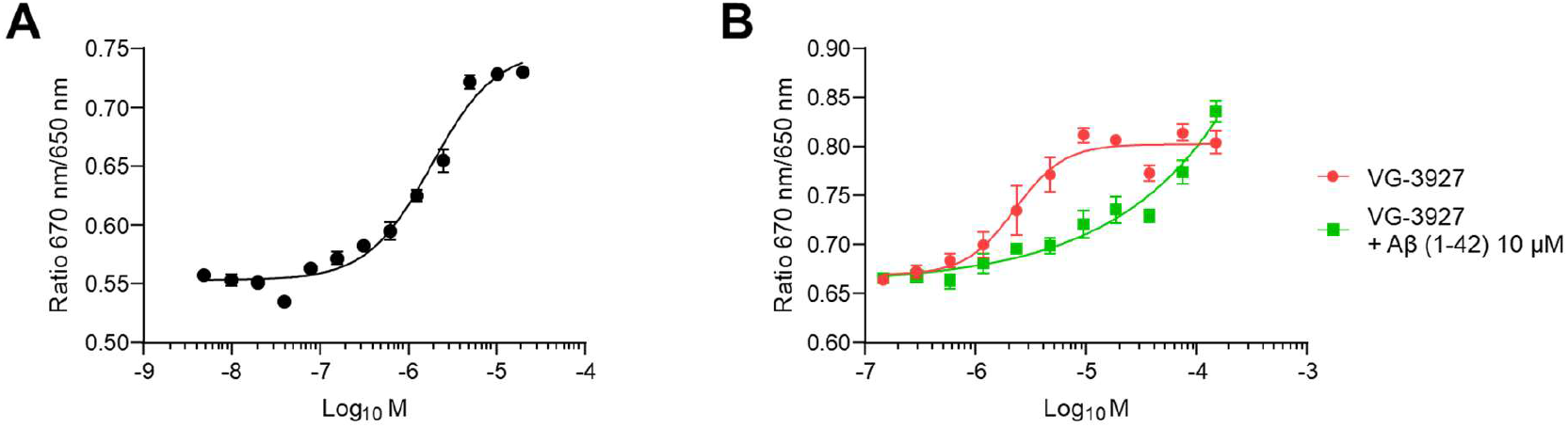
Aβ interferes with VG-3927 for TREM2 binding by MST. (A) MST binding curve showing direct interaction of Aβ_1–42_ with TREM2 under optimized PEG-400 buffer conditions. (B) MST analysis of VG-3927 binding to TREM2 in the absence (red) or presence (green) of 10 μM Aβ_1–42_. Data are mean ± SEM, n = 3.

### 2.4 Aβ Shifts the VG-3927 Dose–Response Curve in the NFAT Reporter Assay

To evaluate functional consequences, we employed a Jurkat TREM2–DAP12 NFAT reporter assay. Aβ itself acts as a partial TREM2 agonist at micromolar concentrations, consistent with reports that Aβ engagement of the hydrophobic site triggers downstream signaling. A sub-maximal concentration of 1 μM Aβ was selected for co-treatment experiments. Co-treatment with 1 μM Aβ and escalating VG-3927 concentrations produced a rightward shift of the VG-3927 dose–response curve (Figure 3), consistent with competitive antagonism. VG-3927 alone yielded an EC^50^ of 88.2 ± 21.5 nM, whereas co-treatment with 1 μM Aβ increased the EC^50^ approximately 4-fold to 351.2 ± 31.9 nM (*P* < 0.01). The preserved maximal efficacy at high VG-3927 concentrations further supports a competitive (rather than non-competitive or allosteric) mechanism, where sufficiently high agonist concentrations can overcome Aβ occupancy.

**Figure 3.**
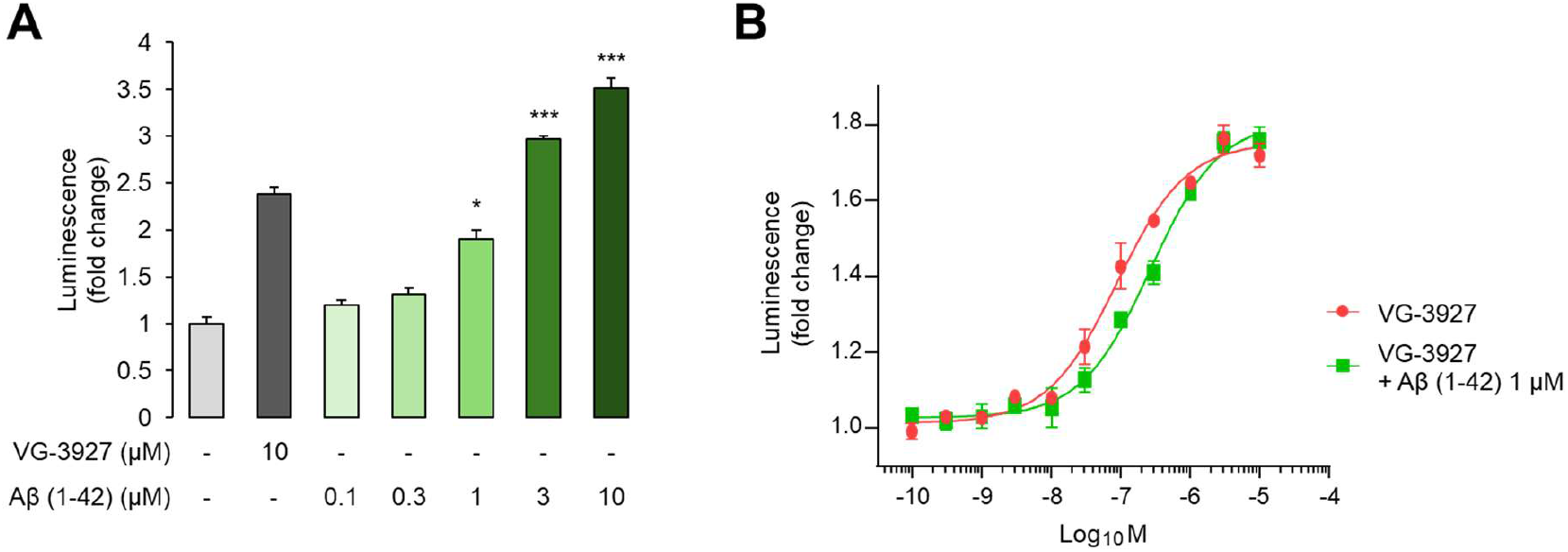
Aβ shifts the VG-3927 dose–response curve in Jurkat TREM2–DAP12 NFAT reporter assay. **(A)** NFAT reporter activity showing Aβ as a TREM2 agonist. 1 µM Aβ (dashed line) was selected for co-treatment. **(B)** VG-3927 dose–response curves alone (red) and with 1 µM Aβ_1–42_ co-treatment (green). Aβ induces a rightward shift of the EC_50_, consistent with competitive antagonism. Co-treatment with Aβ1–42 induced a rightward shift in the VG-3927 dose–response curve, consistent with interference at an overlapping binding site. Data are mean ± SEM, n = 3. *P <0.05, ***P < 0.001 analyzed by Student’s unpaired two-tailed t test.

### 2.5 Aβ Attenuates VG-3927-Induced p-SYK Signaling

To examine whether Aβ-mediated interference with VG-3927 binding affects proximal TREM2 signaling, we measured phosphorylation of spleen tyrosine kinase (SYK), an immediate downstream effector of TREM2–DAP12 activation. Cells were treated with increasing concentrations of VG-3927 in the absence or presence of 1 μM Aβ_1–42_, and p-SYK levels were assessed by western blotting **(Figure 4)**. Densitometric analysis showed that co-treatment with Aβ_1–42_ reduced VG-3927-induced p-SYK signaling. These data are consistent with reduced VG-3927 engagement at the TREM2 ectodomain and attenuation of downstream signaling output.

**Figure 4.**
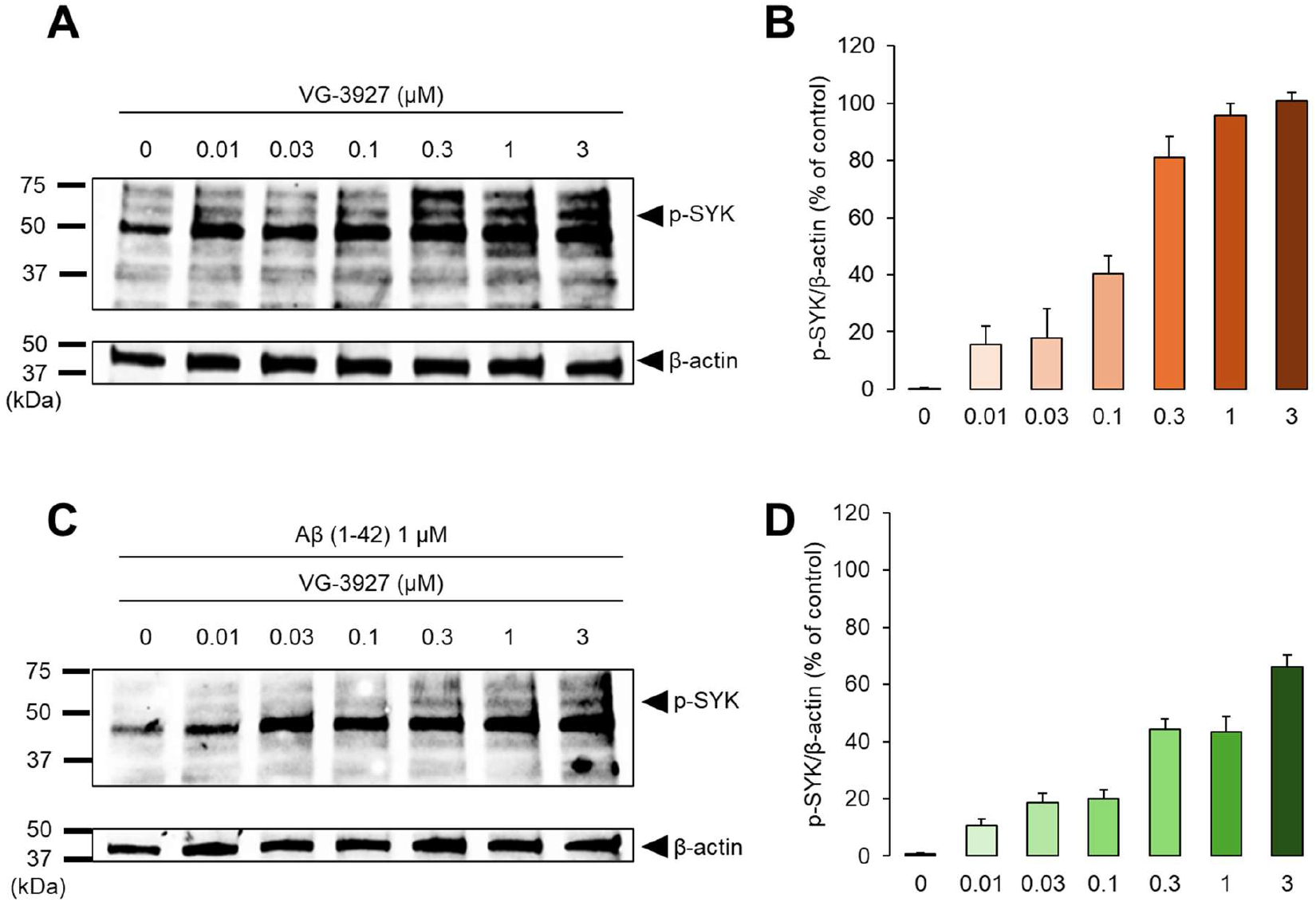
Aβ reduced VG-3927-induced p-SYK signaling. (A) Representative western blots of phospho-SYK (Y525/526) and β-actin in cells treated with increasing concentrations of VG-3927 in the absence of Aβ_1–42_. (B) Densitometric quantification of phospho-SYK normalized to β-actin and expressed relative to the 3 μM VG-3927-alone condition, which was set to 100%. (C) Representative western blots of phospho-SYK (Y525/526) and β-actin in cells treated with increasing concentrations of VG-3927 in the presence of 1 μM Aβ_1–42_. (D) Densitometric quantification showing that co-treatment with Aβ_1–42_ reduced VG-3927-induced p-SYK signaling. Data are presented as mean ± SEM from three independent experiments (n = 3).

## 3. Discussion

In this study, we identify an additional binding mode for VG-3927 at the TREM2 ectodomain hydrophobic groove, a surface that overlaps the Aβ binding pocket. This finding, prompted by the docking observations from our previous study for the VG-3927 analog C1, provides the first experimental evidence that a small-molecule TREM2 agonist, in addition to its established transmembrane mechanism, also engages the same ectodomain surface utilized by Aβ and other endogenous ligands.^23^ While our previous study focused on medicinal chemistry optimization and pharmacokinetic improvement of the VG-3927 scaffold, our work provides experimental validation of an additional ectodomain binding mode for VG-3927 itself and, critically, identifies the specific pocket as the hydrophobic groove (CDR1–CDR3), the same surface used by Aβ for TREM2 engagement. Importantly, our findings do not negate the molecular glue and PAM mechanism proposed by Vigil Neuroscience; rather, they suggest that VG-3927 possesses a dual binding mode, engaging both the transmembrane clustering interface and the ectodomain hydrophobic groove.^14-17^

The overlap between the VG-3927 and Aβ binding sites has several important implications. First, in the amyloid-rich microenvironment of the AD brain, local Aβ concentrations near microglial processes may partially interfere with VG-3927 engagement at the TREM2 ectodomain, attenuating agonist efficacy. Our data showing ∼50% reduction in p-SYK signal at 1 μM Aβ suggests this interference is pharmacologically meaningful but does not completely abolish agonist activity. The observation that VG-3927 retains partial function even in the presence of Aβ is consistent with its dual binding mode: even when Aβ occupies the ectodomain hydrophobic groove, VG-3927 can still promote TREM2 signaling through its molecular glue activity at the transmembrane level. This is also consistent with the developer’s description of VG-3927 as a PAM that synergistically activates TREM2 together with endogenous ligands including Aβ.^22^

The finding that TREM2’s hydrophobic groove serves as a shared binding surface for multiple ligands, including Aβ, apoE, TDP-43, and now VG-3927, is consistent with the “promiscuous binding surface” model proposed by Greven et al.^16^ This shared utilization raises the possibility that endogenous TREM2 ligands may modulate the efficacy of small-molecule agonists in a context-dependent manner. For instance, in brain regions with high apoE deposition, apoE may also interfere with VG-3927 binding at the ectodomain site, adding another layer of complexity to TREM2 pharmacology *in vivo*.

The detergent sensitivity we observed provides a cautionary note for biophysical assay design. Tween-20 is routinely included in MST, SPR, and other binding assays to prevent aggregation and non-specific adsorption.^25^ Our results indicate that for targets with hydrophobic binding pockets, detergent selection must be carefully considered. The successful substitution of PEG-400 for Tween-20 may be broadly applicable to other hydrophobic pocket targets and should be considered in future TREM2 binding studies. Given that our group previously confirmed C1 binding to TREM2 by both MST and SPR, retrospective analysis of detergent conditions in those assays may further clarify the role of the hydrophobic pocket in this chemotype’s binding.^23^

Therapeutically, our findings suggest that combination strategies, pairing TREM2 agonists with Aβ-lowering therapies such as anti-Aβ antibodies (e.g., lecanemab, donanemab), could synergistically enhance TREM2-mediated microglial function by reducing Aβ-mediated ligand occupancy at the ectodomain site.^26, 27^ Additionally, the identification of this second binding site provides a structural foundation for designing next-generation TREM2 agonists that selectively engage one site, avoid Aβ interference, or simultaneously occupy both sites for enhanced potency.

Several limitations should be acknowledged. The DiffDock-L docking, while state-of-the-art, would benefit from validation by co-crystallography or cryo-EM. The MST competition experiments used Aβ at 10 μM, which exceeds typical parenchymal concentrations but may be relevant to plaque-proximal microenvironments. The extent of interference at lower, physiologically relevant Aβ concentrations remains to be quantified. Future studies using iPSC-derived microglia in amyloid-bearing co-culture systems, as well as *in vivo* assessment in TREM2-humanized amyloidosis models, would provide more translational validation.

## 4. Conclusions

Overall, our study supports the presence of an ectodomain hydrophobic-groove interaction mode for VG-3927 and suggests that Aβ can interfere with this interaction, thereby modulating downstream TREM2 signaling. These findings refine the current model of VG-3927 pharmacology and provide mechanistic insight into how amyloid-associated ligand occupancy may influence the activity of TREM2-targeted agonists in Alzheimer’s disease. These observations also suggest that combination strategies pairing TREM2 agonists with Aβ-lowering therapies may help mitigate ligand-mediated interference at the ectodomain, potentially enhancing therapeutic efficacy.

## 5. Materials and Methods

### 5.1 Molecular Docking with DiffDock-L

The TREM2 structure was obtained from the Protein Data Bank (PDB ID: 5ELI).^15^ VG-3927 was prepared as a 3D conformer using RDKit. Blind docking was performed using DiffDock-L with default parameters, generating 40 poses per run.^24^ Poses were clustered by RMSD (cutoff 2.0 Å) and ranked by confidence scores. The top-ranked poses were analyzed for protein–ligand interactions using BIOVIA Discovery Studio Visualizer.

### 5.2 Microscale Thermophoresis (MST)

MST was performed on a Monolith NT.115 instrument (NanoTemper Technologies). Recombinant human TREM2 ectodomain (residues 19–174, BioTechne, Minneapolis, MN, USA) was labeled using the Monolith His-Tag Labeling Kit RED-tris-NTA (2^nd^ Generation). MST buffer consisted of PBS with either 0.05% Tween-20 or 0.05% PEG-400. VG-3927 and Aβ_1-42_ were titrated in 16-point serial dilutions. For competition experiments, TREM2 was pre-incubated with 10 µM Aβ for 15 min at room temperature before VG-3927 titration. Data were analyzed using MO.Affinity Analysis software v2.3. K_d_ values were determined from three independent experiments.

### 5.3 Jurkat TREM2–DAP12 NFAT Reporter Assay

Jurkat cells stably expressing human TREM2, DAP12, and NFAT-luciferase reporter were maintained in RPMI 1640 with 10% FBS. Cells were seeded in 96-well plates (5 × 10^4^ cells/well) and treated with serial dilutions of VG-3927 ± 1 µM Aβ_1–42_ for 12 h. Luciferase activity was measured using Bright-Glo (Promega). EC_50_ values were determined by four-parameter logistic regression in GraphPad Prism 10. For Aβ dose selection, a preliminary dose response confirmed Aβ partial agonism and 1 µM was chosen as a sub-maximal concentration.

### 5.4 Western Blot

Cells were treated with VG-3927 (0, 0.01, 0.03, 0.1, 0.3, 1, 3 µM) ± 1 µM Aβ_1–42_ for 15 min. Lysates were prepared in RIPA buffer with phosphatase and protease inhibitors (Roche). Equal protein amounts (20 µg) were loaded on 4-15% SDS-PAGE gels and transferred to PVDF membranes. Primary antibodies: anti-phospho-SYK (Y525/526, Cell Signaling), β-actin (Cell Signaling). Secondary: HRP-conjugated anti-rabbit IgG. Detection: ECL substrate (Thermo Fisher). Densitometric analysis was performed using ImageJ. Phospho-SYK levels were normalized to β-actin and expressed relative to the 3 μM VG-3927-alone condition, which was set to 100%.

### 5.5 Statistical Analysis

All experiments were performed in ≥ 3 independent biological replicates. Data are mean ± SEM. Statistical comparisons: unpaired two-tailed Student’s *t*-test. All analyses were performed in GraphPad Prism 10.

## Author Contributions

M.T.G. and S.C. designed the study. S.C. performed experiments and analyzed data. M.T.G. supervised the project. All authors wrote the manuscript and approved the final version.

## Notes

The authors declare no competing financial interest.

## Insert Table of Contents artwork here

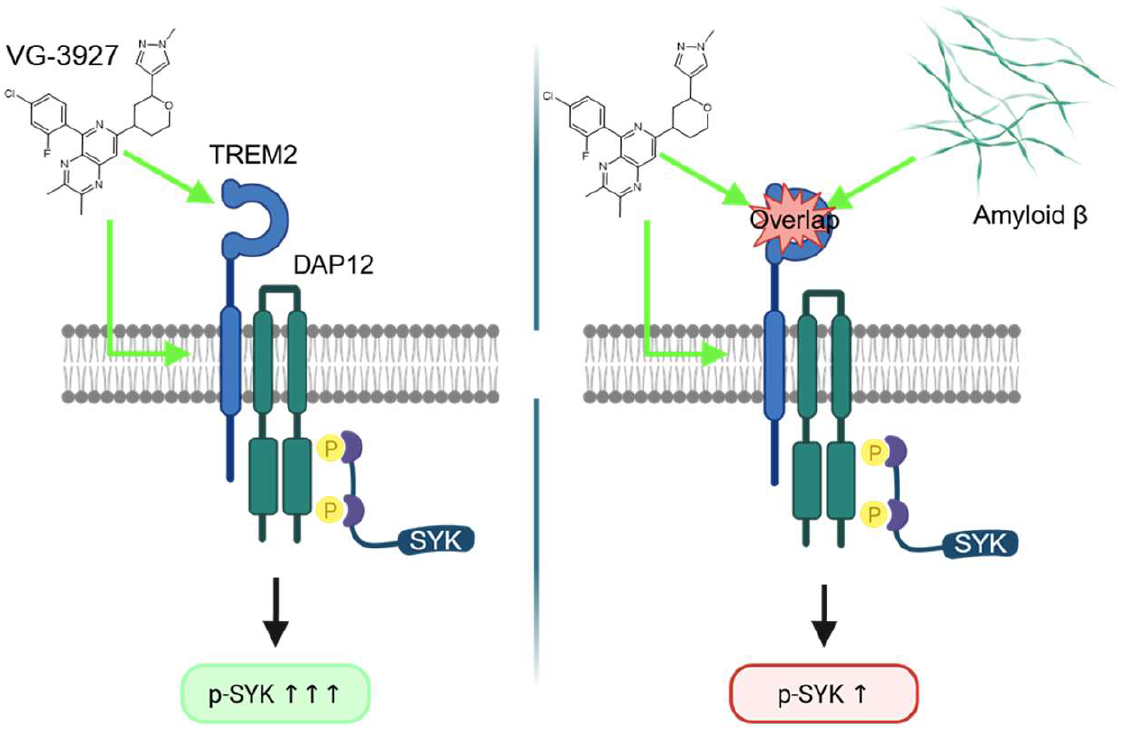

